# Mineral fertilization reduces the drought resistance of soil multifunctionality in a mountain grassland system through plant-soil interactions

**DOI:** 10.1101/2024.09.19.613911

**Authors:** Gabin Piton, Arnaud Foulquier, Lionel Bernard, Aurélie Bonin, Thomas Pommier, Sandra Lavorel, Roberto Geremia, Jean Christophe Clement

## Abstract

Increasing droughts threaten soil microbial communities and the multiple functions they control in agricultural soils. These soils are often fertilized with mineral nutrients, but it remains unclear how this fertilization may alter the capacity of soil multifunctionality (SMF) to be maintained under drought, and how plant-soil interactions shape these effects. In this study, we used a mountain grassland soil to test the interactive effect of mineral nutrient (Nitrogen and Phosphorous) addition and drought on SMF with and without plants (*Lolium perenne*) in a mesocosm experiment. We calculated SMF based on 8 microbial properties associated with the capacity of soil microbes to store carbon (C), nitrogen (N) and phosphorous (P) in their biomass, and to process these elements through organic matter depolymerization, mineralization, nitrification and denitrification processes. To investigate mechanisms underlying the SMF response we characterized the associated changes in soil stoichiometry and microbial community composition using 16S and 18S rRNA amplicon sequencing. Our results showed that fertilization decreased the SMF drought resistance when plants were present, but the opposite was observed in the unplanted mountain grassland soil. Our analysis suggested this was due to the interaction of plants, fertilization and drought in influencing four coupled properties related to high SMF: high soil moisture, low microbial C limitation, high bacterial diversity and low bacteria gram positive:gram negative ratio. Altogether, our results suggested that reducing the use of mineral fertilizer for plant production in mountain grassland could improve the ability of their soils to maintain their multifunctionality during drought period. Finally, our study clearly further demonstrated the importance of plant in the complex responses of SMF to global changes and showed that combining stoichiometric and microbial diversity assessment represents a powerful approach to disentangle the underlying mechanisms.

## 1 Introduction

Soil microbial communities regulate key processes in terrestrial biogeochemical cycles, underlying critical ecosystem services such as soil fertility maintenance, and climate change mitigation through carbon sequestration (Crowther et al. 2019). The concept of ecosystem multifunctionality has been developed to integrate multiple ecosystem functions into composite indices, providing a global assessment of ecosystem performance (Zavaleta et al. 2010, Maestre et al. 2012). Soil multifunctionality (SMF) specifically refers to the soil’s ability to maintain multiple functions simultaneously, including carbon and nutrients storage and recycling (Garland et al. 2021), with most of these functions being controlled by soil microbial communities (Delgado-Baquerizo et al. 2016b).

With current global change, it becomes essential to understand the mechanisms controlling microbial community properties underlying SMF. Microbial processes in soil are strongly influenced by abiotic factors such as pH, humidity, redox potential and temperature (Fierer 2017). Microbial metabolism also depends on the availability and composition of nutritional resources, often investigated through ecological stoichiometry, a framework that assesses chemical elements balance to predict nutrient storage and fluxes among different biotic and abiotic pools (Zechmeister-Boltenstern et al. 2015). Stoichiometric theory predicts that soil microorganisms immobilize and store the most growth-limiting elements, while excess elements are mineralized and released in the environment. However, some specific microbial functions, including nitrification, denitrification or depolymerization of specific substrates also depend on the presence of specific microbes or microbial consortia (Crowther et al. 2019). Consequently, a high diversity of microbes is expected to support multiple soil functions. This is supported by observations linking higher microbial diversity and specific community composition to a high magnitude and stability of SMF (Delgado-Baquerizo et al. 2016b, 2017, Maron et al. 2018), while functional redundancy is expected to weaken this relationship in some cases (Allison and Martiny 2008, Delgado-Baquerizo et al. 2016a, Trivedi et al. 2019, Yang et al. 2024). Altogether, this indicates that maximizing SMF requires specific abiotic and stoichiometric conditions along with a diverse microbial community able to sustain multiple functions.

Grasslands cover 69% of the global agricultural land area and their management has been drastically intensified in many part of the world over the last century, particularly through the increased use of mineral fertilizers (Bardgett et al. 2021). On the other hand, drought intensity and frequency have increased in many regions, with dramatic consequences on ecosystem functioning (Shukla et al. 2019). While SMF is of central importance for grassland ecosystem services (Grigulis et al. 2013, Garland et al. 2021), our understanding of how fertilization influences the drought resistance of grassland SMF is still limited. Life history strategy theory proposes that r-strategists (ie. copiotrophic) microbes thrive in resource-rich environments with high C and nutrients availability and exhibit a low resistance but a high resilience to climatic stress (De Vries and Shade 2013). Stoichiometric theory further suggests that the rapid growth of r-strategists is dependent on a balanced supply of resources to meet their biomass needs (Mooshammer et al. 2014). Taking into account the tight link between the nutritional and water stress niches of soil microbes (Fierer 2017, Piton et al. 2023, Camenzind et al. 2024), fertilization may improve resource availability and/or their stoichiometric balance, which may also select for microbes with lower resistance to subsequent drought (Piton et al. 2020c). However, such a negative effect of nutrient enrichment through selection of r-strategists taxa could be balanced by potential physiological benefits of high nutrient availability for microbes under conditions of osmotic stress. Indeed, fertilizer nutrients remain in the soil at the beginning of drought (ie. not assimilated by plants, microbes or lost from soil) and may sustain the increased bacterial N-demand for osmolyte production or other stress resistance mechanisms (Schimel et al. 2007).

Beyond life history strategies and microbial physiology, microbial diversity play also a key role in ecosystem resistance. Nutrient addition to soil has been shown to decrease bacterial and fungal richness and alter their community composition (Zhou et al. 2020), potentially decreasing the resistance of ecosystem functioning to climatic stress. Whether this microbial diversity effect is driven by microbial alpha diversity (e.g. richness) or by changes in community composition and life history strategy remains debated (Yachi and Loreau 1999, Tardy et al. 2014, Yang et al. 2022, Osburn et al. 2023). Increases in the gram positive : gram negative (GP:GN) and the fungal : bacterial (F:B) ratios represent one of the most consistent shifts of microbial community composition under drought. This has been mainly attributed to the properties of gram positive and fungal cell walls, and a dominance of K-strategists and stress tolerance traits within these groups (De Vries and Shade 2013, Canarini et al. 2017, Naylor and Coleman-Derr 2018). Such consistent patterns suggest that factors changing the GP:GN or F:B ratio, such as C limitation (Fanin et al. 2018) and mineral fertilization (Bardgett et al. 1996, De Vries and Bardgett 2012), could have important consequences for microbial community stability and its response to environmental stress (De Vries and Shade 2013).

Plants also play a central role in mediating the effects of fertilization on soil microbial communities, as their growth depend on nutrient and water uptake from the soil. Plants growth is also associated with the release of C-rich molecules from rhizodeposition and tissues senescence (Bardgett et al. 2008). Experiments support the central role of C, nutrient and water dynamic in linking plant and microbial responses to nutrient addition and drought (Bloor et al. 2018, De Vries et al. 2018, Piton et al. 2020a, 2020c). However, most studies examining the interactions between drought and nutrient availability on SMF have been conducted under laboratory conditions in the absence of plant (Luo et al. 2019, Dong et al. 2022) or have focused on plant species and fertilization rates that are not representative of managed grasslands (Preece et al. 2020). To better understand how fertilization influences SMF response to drought in grasslands, new plant-soil experiments combining these factors are needed.

This study aimed at investigating how mineral fertilization and drought interact to impact the SMF of a mountain grassland. To assess the role of plants in mediating these effects, we tested the fertilization effect in mesocosms with and without plants. Mineral fertilization increases the level of nutrients in soil and, when plant are present, stimulates their growth and the associated input of fresh C to the soil, balancing microbial resource stoichiometry. Based on life history strategy and stoichiometric theories, we thus hypothesized that such effects on soil resource levels and stoichiometric balance, could alter soil microbial diversity with a selection of a microbial communities with a lower capacity to maintain SMF under drought conditions.

## 2 Materials and methods

### 2.1 SOIL SAMPLING

Soil was collected in a grassland localized in the Chartreuse mountain range of French Alps (45°36’ N, 5°90’E), at 930 m a.s.l. with a mean annual temperature of 8.1°C and 1049mm of mean annual precipitation. Management was extensive since decades with one mowing per year and no fertilization. The 15 first cm of the topsoil (3.6% of C and 0.4% of N) were extracted in spring (5^th^ and 6^th^ of April 2018), and sieved at 7mm to remove plant residues. After homogenization, soil was used to fill 32 mesocosms (25cm diameter and 40cm depth). All mesocosms were transported in the alpine green-house of the Jardin du Lautaret (Grenoble), with an air temperature maintained at 20°C during the experiment, approximating the study site’s daily mean temperature during the growing season. Mesocosms were weighted prior the start of the experiment (mean fresh soil weight: 11.8 kg, 95% confidence interval ± 0.1kg). All along the experiment, control mesocosms (unplanted soil, control climate) were weighted twice a week and compared with their initial weight to assess water loss. All mesocosms received the same amount of water equal to this water loss in control mesocosms (in average 300mL per week), to maintain a stable soil moisture in mesocosms equal to the initial field value (0.24 g water per g of fresh soil, 66% of maximum water holding capacity). During the first 30 days, soil water content was kept constant for all mesocosms. In parallel, seeds of *Lolium perenne* from Arbiotech® (Saint Gilles, France) were germinated in individual pots filled with the same soil used in the mesocosms until seedling establishment. *L. perenne* was used as a phytometer because it is usually sown in managed grasslands in the French Alps (Loucougaray et al. 2015, Legay et al. 2017). After 30 days of growth (2-3 leaves stage), 25 individuals of *L. perenne* were planted in 16 mesocosms, with a 4 cm distance between each individual and with mesocosm edge. At day 45 (tillering start of *L. perenne*), half of the mesocosms (planted and unplanted soil) were fertilized with mineral N (50%-N as NO_3_^-^ and 50%-N as NH_4_^+^) and P (P_2_O_5_ equivalent to 150 kg N/ha and 80kg P_2_O_5_/ha representing the regional recommendation for nutrient addition in intensive sown grasslands (Guide régional de fertilisation 2016, Chambre d’Agriculture). Twenty days after fertilization (day 65), watering of 16 mesocosms (4 unplanted soil/unfertilized, 4 unplanted soil/fertilized, 4 planted soil/unfertilized, 4 planted soil/fertilized) was stopped. After 32 days without water addition for the drought treatment (day 97 of the experiment), corresponding to a moderate drought simulation in mountain grasslands (Karlowsky et al. 2018, Brunner et al. 2022, Möhl et al. 2023), three soil cores (5cm diameter, 15cm depth) were sampled in the center of each mesocosm on the location of three *L. perenne* individuals. Soil cores were then pooled in a composite sample to assess plant biomass and soil stoichiometry, microbial community biomass, activity and diversity.

### 2.2 Plant analyses

Plants shoots and roots in planted mesocosms were manually sorted from the three soil cores. Plants shoots and roots were separated and dried at 70°C for a week and weighed to measure shoots and roots biomass. All plant material was ground and analyzed for C and N contents using an elemental analyzer (FlashEA 1112: Fisher 302 Scientific, Waltham, Massachusetts, USA).

### 2.3 Soil analyses

Composite soil samples were split in subsamples, some of which were immediately frozen in liquid nitrogen and stored at -80°C until molecular analyses. Subsamples for enzymatic analyses were frozen at -20°C until analysis. Subsamples for C, N and P pools were stored at 4°C and analyzed within 24h.

Subsamples of 5g of fresh soil were oven-dried at 70°C for 1 week and weighed to determine soil water content (SWC) (Robertson et al. 1999). Soil subsamples were air dried and ground to powder to measure total C and N contents using a FlashEA 1112 elemental analyzer (FlashEA 1112: Fisher 302 Scientific, Waltham, Massachusetts, USA), and to determine soil pH in a 1:2.5 (soil:distilled water) solution. Dissolved organic carbon (DOC), nitrate (NO_3_^-^), ammonium (NH_4_^+^), total dissolved nitrogen (TDN) were extracted on 10 g of fresh soil using 0.5M K_2_SO_4_ solution following Jones and Willett (2006). DOC concentration was measured on a Shimadzu TOC-L analyzer (Shimadzu Inc.). N and P concentrations were measured on an automated photometric analyzer using standard colorimetric methods (Gallery Plus: Thermo Fisher Scientific, Waltham, Massachusetts, USA). Dissolved organic nitrogen (DON) was calculated as the difference between TDN and total inorganic N (N-NO_3_^-^+ N-NH_4_^+^).PO_4_ was extracted on 10g of fresh soil using 0.5M NaHCO_3_ solution as described by (Brookes et al. 1982) and quantified with method of Murphy and Riley (1962)

### 2.4 Microbial biomass and activities

Microbial biomass carbon (MBC), nitrogen (MBN) and phosphorus (MBP) contents were measured on 10-g of fresh soil, using chloroform fumigation extraction-methods as described by (Vance et al. 1987) for MBC and MBN, and by Brooks et al. (1981) for MBP. A spiking procedure (25µgP-PO_4_/g soil were added on two samples) was used to estimate and correct for P recovery during extraction (Brookes et al. 1982). Microbial biomass element contents were calculated as the difference between fumigated and non-fumigated samples, and adjusted using conversion factors of 0.45, 0.45 and 0.40 for MBC, MBN and MBP respectively (Jenkinson et al. 2004). Indicators of the stoichiometric imbalances between microbial biomass and their resources were calculated as the ratios of resource A:B in the soil solution over microbial biomass A:B (Mooshammer et al. 2014), with A:B being C:N, C:P or N:P ratios. Assuming that the microbial biomass elemental composition represent its nutritional need to grow, an increase of such indicator above 1 indicate that the element B become more limiting over A, in other words its availability relative to A become too low to build new biomass. Values below 1 indicate the opposite situation.

The potential activities of 7 extracellular enzymes (EEA) involved in the decomposition of C-rich substrates (α-Glucosidase, β-1,4-Glucosidase, β-D-Cellobiosidase, and β-Xylosidase), N-rich substrates (β-1,4-N-acetylglucosaminidase and leucine aminopeptidase) and P-rich substrates (phosphatase) were estimated using standardized fluorimetric techniques (Bell et al. 2013). Briefly, 2.75-g of soil were homogenized (1-min in a Waring blender) in 200-ml of sodium acetate buffer solution adjusted at soil pH (5.8). The soil slurries were added in duplicate to 96-deep-well microplates followed by the addition of a substrate solution for each enzyme at enzyme saturation concentration. Duplicated standard curves (0-100-µM concentration) were prepared by mixing 800-ml of soil slurry with 200-ml of 4-methylumbelliferone (MUB) or 7-amino-4-methylcoumarin (MUC) in 96-deep-well microplates for each soil sample. Microplates were incubated during 3-h (dark, 175-rpm, 20°C), and centrifuged at 2900-g for 3-min. Then soil slurries (250-µL) were transferred into black Greiner flat-bottomed microplate and scanned on a Varioskan Flash reader (Thermo Scientific) using excitation at 365-nm and emission at 450-nm (Bell et al. 2013). After correcting for negative controls, potential enzyme activities were expressed as nmol g soil^-1^ h^-1^. The activities of the seven enzymes were summed to provide a measure of cumulative extracellular enzyme activity (EEA).

Potential N mineralization (PNM) was estimated after incubation of 10-g of fresh soil under anaerobic conditions for 7 days at 40°C in the dark (Wienhold 2007), inducing accumulation of mineralized NH_4_^+^. PNM rate was calculated as the difference between NH_4_^+^ content before and after incubation (µgN/g dry soil/day). Potential nitrification enzyme activity (NEA) was estimated according to (Koper et al. 2010) following (Dassonville et al. 2011). Briefly, 3 g of each soil were incubated under aerobic conditions (180 rpm, 28°C, 10 h) in a solution of 50µg N-(NH4)2SO4/g dry soil and rates of NO_2_ and NO_3_ production were measured after 2, 4, 8 and 10 hr by ionic chromatography (DX120; Dionex, Salt Lake City, UT, USA). Maximal nitrification rate (NEA) was assessed by plotting nitrification rates along the gradient of NH_4_–N concentrations (Lineweaver and Burk 1934). Potential denitrification activity (DEA) was estimated following Attard et al. (2011). Briefly, 10 g dry weight (dw) soil were placed at 28°C under anaerobic conditions using 90:10 He:C_2_H_2_ mixture inhibiting N_2_O-reductase activity. Each sample was supplemented with 3 ml KNO_3_ (50 µg N–NO_3_ g^−1^dw), glucose (0.5 g C/g dw) and sodium glutamate (0.5 g C/g dw) (Attard et al. 2011), completed with distilled water to reach water holding capacity. N_2_O was measured at 2, 3, 4, 5 and 6 hr using a gas chromatograph (microGC R3000; SRA Instruments, Marcy l’Etoile, France). Substrate-induced respiration (SIR) was measured following (Anderson and Domsch 1978). Briefly, 5 g of soil samples were placed in airtight flask, with 1.2 mg of C-glucose/g dry soil, completed with distilled water to reach water holding capacity. Then, samples were incubated at 28°C during 5 hours with CO_2_ concentrations measured each hour. The slope of the linear regression between time and with CO_2_ concentrations was used to estimate aerobic respiration (g C-CO_2_ ^−1^ h^−1^).

### 2.5 Soil multifunctionality

A soil multifunctionality (SMF) index was calculated using the widely used method presented by Maestre et al. (2012). Eight microbial properties and functions (MBC, MBN, MBP, SIR, EEA, PNM, NEA, DEA) were first standardized using Z-score transformation and then averaged. These variables were selected because they capture the capacity of soil microbes to store C, N and P in their biomass and to process these elements through organic matter depolymerisation, mineralization and downward transformations (nitrification and denitrification).

### 2.6 Molecular analyses

Soil microbial diversity was estimated using environmental DNA metabarcoding targeting two universal DNA markers (Table S1), one amplifying all Bacteria (v4 region of the 16S rRNA gene) and one amplifying all Eukaryota (v7 region of the 18S rRNA gene). Extracellular DNA was extracted from 10g of soil using the phosphate buffer procedure described in (Taberlet et al. 2012). PCR amplifications were performed in a 20-μL volume containing 10 μL of AmpliTaq Gold 360 Master Mix (Applied Biosystems, Foster City, CA, USA), 0.2 μM of each primer, 0.16 μL of 20 mg.ml-1 bovine serum albumin (BSA; Roche Diagnostics, Basel, Switzerland), and 2 μL of 1/10 diluted DNA extract. Thermocycling conditions (Table S1) followed recommendation from Taberlet et al. (2018) and Guardiola et al. (2015). The forward and reverse primers were tagged with unique combinations of eight-nucleotide labels. PCR products were purified using the MinElute™ PCR purification kit (Qiagen, Hilden, Germany) and mixed in equal volumes. Library preparation and sequencing were performed at Fasteris (Fasteris SA, Geneva, Switzerland) using the MetaFast protocol (www.fasteris.com/metafast). The bacterial and eukaryote libraries were sequenced on an Illumina MiSeq (2 x 250 bp paired-end reads) and HiSeq 2500 ((2×150 bp paired-end reads) platforms, respectively. Potential contaminations were tracked by including negative controls at the extraction and PCR steps. Tag jumps caused by chimeras were accounted for at the bioinformatic level by monitoring unused tag combinations (Schnell et al. 2015).

Sequencing data were then analysed using the OBITools software package (Boyer et al. 2016) and R scripts following (Zinger et al. 2019). Firstly, paired-end reads were assembled and then assigned to the corresponding PCR replicate on the basis of primer sequences (two mismatches allowed) and tag combinations (no mismatches allowed). Secondly, reads were dereplicated. Low quality sequences (i.e. sequences containing “Ns”), sequences observed less than 100 times, and sequences whose length fell outside the expected length range (<45bp and <36bp for the bacterial and eukaryote markers, respectively) were discarded (Taberlet et al. 2018). Thirdly, unique sequences were clustered at 97% similarity into operational taxonomic units (OTUs) using the Sumaclust package (Mercier et al. 2013). Only the cluster centres were kept for taxonomic annotation with the ecotag program from the OBITools package (Boyer et al. 2016). OTUs whose similarity score of the best match (best identity score from ecotage program) in the reference database was below 0.95% were excluded. Then, potential contaminations were removed by excluding all OTUs whose absolute number of reads was highest in negatives control. PCR replicates were summed for each sample and rarefied to the same sequencing depth equal to the lowest number of sequences observed in a sample. Rarefaction curves were also obtained for each sample using the *vegan* R package (Oksanen et al. 2011) to assess to which extent the sequencing depth captured the diversity present in our samples (Figure S1). All fungal sequences were extracted from the Eukaryota dataset to narrow down the subsequent analyses to bacterial and fungal communities. The gram positive to gram negative ratio of soil bacteria was calculated using relative abundance of 16S rDNA OTUs as in Orwin et al. (2018), which relied on the Gupta (2011) classification of bacteria phylum based on the presence of the Hsp60 insert in their genomes (Gram positive : Actinobacteria, Firmicutes, Deinococcus-Thermus and Chloroflexi ; Gram negative : Proteobacteria, Acidobacteria, Bacteroidetes, Chlamydiae, Gemmatimonadetes, Nitrospirae, Planctomycetes, Spirochaetes, Verrucomicrobia, Armatimonadetes, Elusimicrobia and Fusobacteria).

### 2.7 Statistical analyses

Three-way analysis of variance (ANOVA) were used to test the effect of plant presence, fertilization, drought, and their interactions on microbial and soil properties. Two-way ANOVA was used to test the effect of fertilization, drought and their interaction on total plant biomass. Model residuals were tested for normality and homoscedasticity and variables were log-transformed when necessary. Three-way permutational multivariate analysis of variance (PERMANOVA) using Bray-Curtis distance matrices were used to test the treatment effects on bacterial and fungal community composition. Microbial community composition data were transformed using Hellinger transformation before PERMANOVA analyses. Significance was set to P < 0.05.

## 3 Results

Significant interaction between drought, fertilization and plant presence were observed for SMF (Table S2), indicating that SMF response to drought changed across the different combinations of fertilization and plant presence treatments (Figure 1). Indeed, drought decreased SMF in unplanted mountain grassland soil only in the absence of fertilization, indicating a lower resistance of SMF. Conversely, when plants were present, SMF did not decrease with drought in unfertilized treatment but did decrease in the fertilized one. Across all treatments, the lowest SMF was observed in the presence of plants, when drought and fertilization were combined. When analyzing individual functions, significant interactions between drought, fertilization and plant presence were also detected (Table S2, and Figure S2). The lower resistance observed in unplanted soil without fertilization was triggered by a decrease of MBC, SIR and MBP, whereas unplanted fertilized soil did not show this negative drought effect, instead positive responses to drought were observed for PNM and MBN. However, this higher resistance associated with fertilization in unplanted mountain grassland soil was not associated with significantly higher value of microbial functions for fertilized soils compared to unfertilized soils under drought (excepted for MBN) because in most cases fertilization has already altered microbial functions in the same direction as drought did (Figure S2). In the presence of plant, drought did not affect microbial functions in the unfertilized treatment, whereas it induced a decrease of MBC, SIR, MBN and PNM when plant had been fertilized. Fertilization and drought also affected plant biomass with an average increase of 220% with fertilization, and a decrease by 30% with drought (Figure S3).

**Figure 1.**
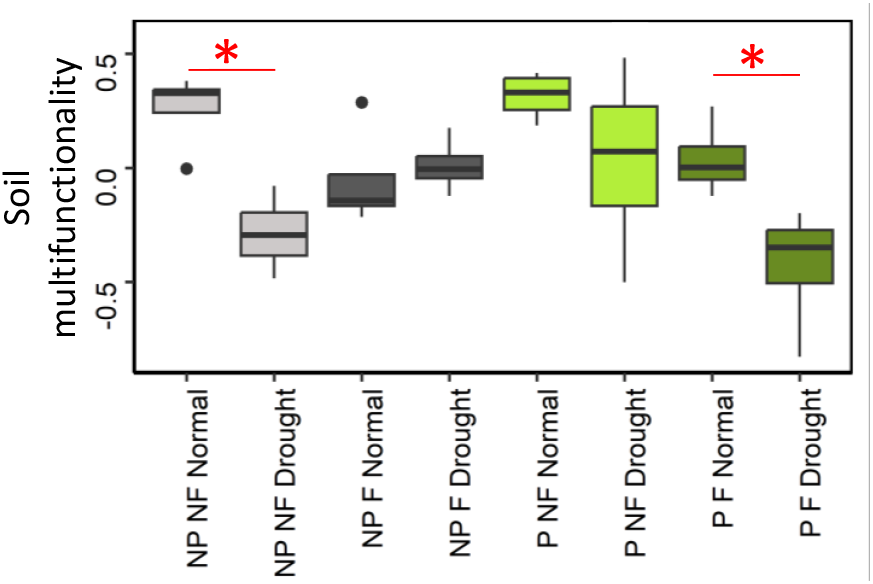
Soil multifunctionality response to drought and fertilisation, with or without plants. Star indicate significant difference between drought and normal climate treatment. NP=No plant, P=Plant presence, F=Fertilized, NF=No fertilization. Grey = unplanted soil, green = planted soil, light colour = unfertilized soil, dark colour = fertilized soil.

Metabarcoding analyses identified 3,181 bacterial OTUs and 624 fungal OTUs across all treatments. Drought significantly decreased bacterial Shannon index but increased fungal richness (Table S2, Figure S4). Bacterial alpha-diversity (Shannon index and OTU richness) also decreased with fertilisation but only in the presence of plants, and fungi showed the lowest richness values when fertilization, plant presence and normal climate were combined (Table S2, Figure S4). Plant presence, drought and fertilisation significantly influenced the composition of bacterial and fungal communities. According to the PERMANOVA models (Table 1), these experimental treatments explained a total of 43% and 39% of the variation in bacterial and fungal community composition, respectively. No significant interaction was observed for fungal community composition, whereas interaction between fertilisation and plant presence was significant for bacteria. Indeed, posthoc tests showed significant effect of fertilization on bacterial community composition in planted soil (p<0.001) but not in unplanted soil (p=0.12).

**Table 1.** P-values and R^2^ associated with the effects of Drought, Fertilization, Plant presence and their interactions on bacterial and fungal community composition. Effects were assessed by PERMANOVA on a Bray-Curtis distance matrix based on bacterial (16S rRNA) and fungal (18S rRNA) operational taxonomic units (OTUs).

| | Drought<br>(D) | Fertilization<br>(F) | Plant<br>presence<br>(P) | D × F | D × P | F × P | D × F × P | Total $R^2$ |
| --- | --- | --- | --- | --- | --- | --- | --- | --- |
| **Bacteria** |  |  |  |  |  |  |  |  |
| p-value | **p<001** | **p<001** | **p<001** | 0.07 | 0.10 | **0.03** | 0.15 |  |
| $R^2$ | **0.10** | **0.12** | **0.07** | 0.03 | 0.03 | **0.04** | 0.03 | 0.43 |
| **Fungi** |  |  |  |  |  |  |  |  |
| p-value | **p<001** | **0.02** | **p<001** | 0.60 | 0.33 | 0.38 | 0.30 |  |
| $R^2$ | **0.10** | **0.05** | **0.13** | 0.02 | 0.03 | 0.03 | 0.03 | 0.39 |

While SMF showed contrasting response to drought and fertilization depending on plants presence, SMF showed consistent relationships with some soil and microbial properties across planting treatments (Figure S5 and S6). In both planted and unplanted soils, SMF was positively correlated with soil moisture, C:N imbalance and bacterial diversity, and negatively correlated with the bacterial gram positive : gram negative ratio (Figure 2). These 4 variables were selected for their consistent relationships with SMF, and variance partitioning showed that 25% was commonly explained by them (Figure S7). Microbial properties explained 24 additional %, with 8 % explained by both microbial properties and 16% explained only by the GP:GN ratio.

**Figure 2.**
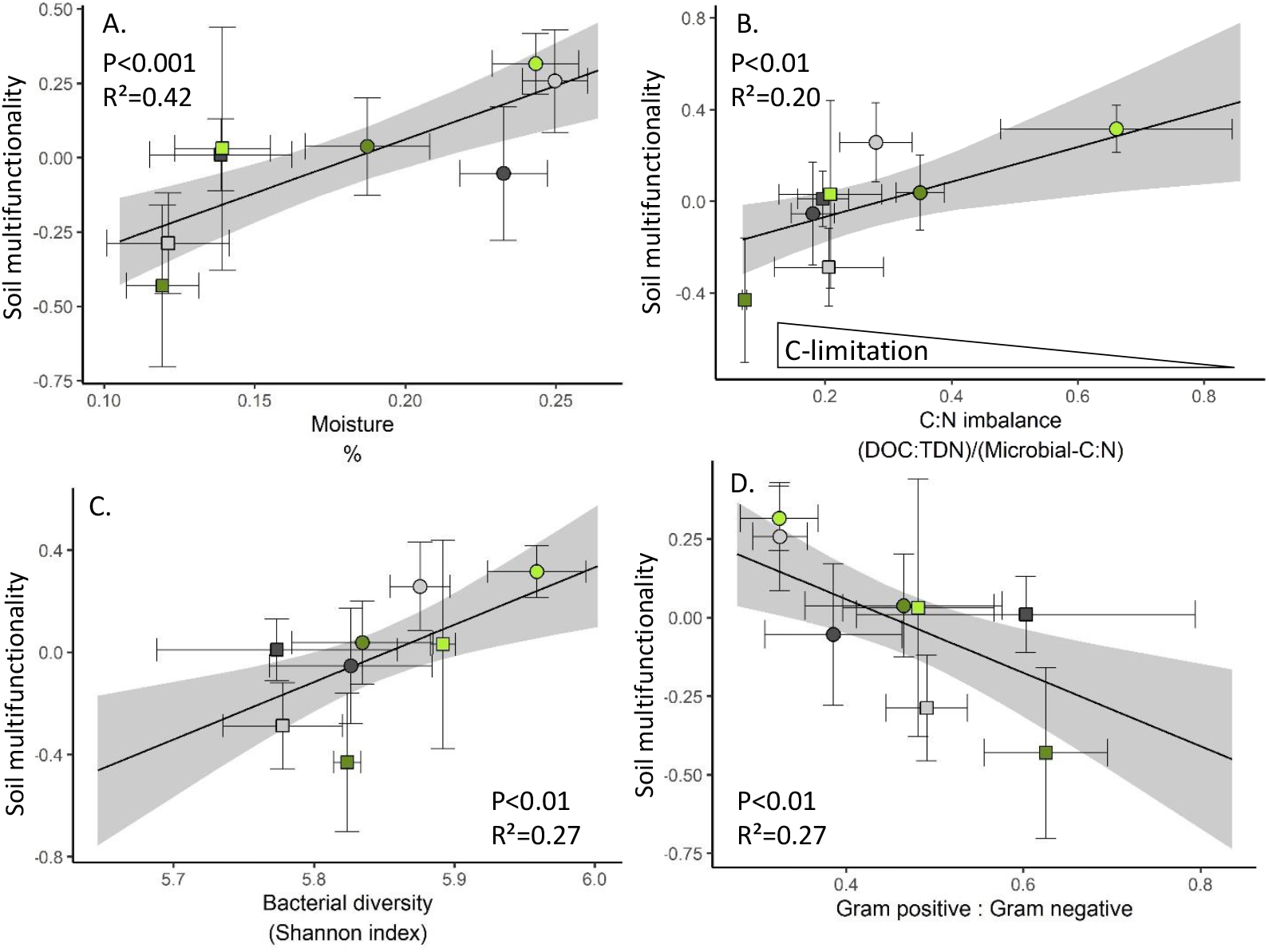
Correlations between SMF and soil moisture (A), C:N imbalance (B.), bacterial alpha diversity (C) and Gram positive : Gram negative ratio (D). Lines represent predicted value and shades represent 95% confidence interval from regression model using all observations (n=32). Points represent mean of each treatments. Error bar around points are 95% confidence interval (n=4). Grey points = unplanted soil, green points = planted soil, light colour points = unfertilized soil, dark colour points =fertilized soil, circle=control climate, square=drought.

## 4 Discussion

Our experiment tested the interactions between fertilization, drought and plant presence on the multifunctionality of a mountain grassland soil. Based on life history strategy theory, we hypothesized that mineral fertilization would reduce the resistance of soil multifunctionality (SMF) to drought. Our results support this hypothesis in planted soils but revealed the opposite in unplanted soils. We propose that these differential responses could be explained by the effect of plant on soil resource stoichiometry balance and microbial diversity.

The role of plants in mediating the interacting effects of drought and fertilization on SMF remains largely unexplored in a unique experiment. Using laboratory microcosms without plants, Luo et al. (2019) found a positive effect of mineral fertilization on the resistance of SMF (indices based on 7 enzyme activities) to dry/rewetting cycles of a cropland soil, consistent with our results in the unplanted grassland soils. However, Dong et al. (2022) tested the effect of an induced N-deficiency on microcosms from grassland soils and found no impact on drought resistance. On the other hand, Eskelinen et al. (2020) studied the interaction between fertilization and watering on plants and soil of a Mediterranean grassland, and observed that increased amount of soil nutrients under fertilization and water stress (no watering), had no effect on microbial biomass and processes. This suggests that the negative effect of fertilization on SMF resistance to drought that we observed in planted soil from a mountain grasslands might not be a generic pattern and likely depends on specific environmental conditions. Supporting this idea, Preece et al. (2020) showed that the significance of the interaction between nutrient input and drought on microbial respiration and metabolic profile in a Mediterranean forested soil was depend on tree-species identity. Our results also indicate that plant can improve the drought resistance of SMF in unfertilized conditions, contrasting with other studies reporting negative effect of plant presence for soil microbial community composition and respiration resistance to drought (Orwin and Wardle 2005, Koyama et al. 2017). However, our study focused on the resistance of SMF to a single, short-term drought event (32 days without water input), while differences of microbial functions between treatments could be attenuated or accentuated with drying time (Borken and Matzner 2009) and repeated droughts can lead to microbial adaptation to such disturbance (Evans and Wallenstein 2014). Additionally, we did not consider the resilience of SMF although it is an important parameter to understand the long term consequences of a disturbance (Wardle and Jonsson 2014). Trade-off between resistance and resilience to drought of microbial communities have been shown across management intensity gradients in grasslands (De Vries et al. 2012, Karlowsky et al. 2018, Piton et al. 2020a, 2020c), suggesting that treatments inducing low resistance of SMF in our study could be associated with better resilience. Our study provides new insights into how fertilization drives SMF resistance to drought in mountain grasslands. However, further research is needed to fully examine the consequences of such interaction between global change factors under different environmental conditions and repeated drought cycles.

### Coupled changes of soil moisture, stoichiometry and bacterial community explain soil SMF responses

To better understand how fertilization can shape the resistance of SMF to drought, it is essential to identify the underlying mechanisms. We hypothesized that changes in resource stoichiometry and microbial diversity in a mountain grassland soil could drive the response of SMF to interacting factors, including drought, nutrient addition and plant presence. Indeed, whether plants were present or not, we observed a consistent link between high SMF and four coupled properties: high soil moisture, C:N imbalance closer to 1 (indicating lower microbial C limitation), high bacterial diversity and low bacteria gram positive : gram negative ratio. These links with soil moisture and stoichiometry confirmed the importance of microbial growth favorable conditions to maximize their multifunctionality, as predicted by stoichiometric and ecohydrological models of microbial biomass and processes (Manzoni et al. 2019). The positive link with bacterial diversity, and the higher diversity observed in the planted mountain grassland soil without fertilization depicting high SMF resistance, support increasing evidence that a diverse microbial community might also be key to maintain high SMF in planted ecosystems exposed to global changes (Delgado-Baquerizo et al. 2016b). Our results are also consistent with studies indicating stronger relationship between SMF and bacterial diversity rather than with fungal diversity (Jia et al. 2022, 2024). SMF also increased with gram positive : gram negative ratio, a trend highly associated with an increase of *Actinobacteria*, a gram positive phylum, and with a decrease of *Proteobacteria*, a gram negative phylum. Abundances of gram negative bacteria, such as *Proteobacteria*, are known to decrease with C limitation (Fierer et al. 2007, Fanin et al. 2018), and water stress (Naylor and Coleman-Derr 2018), as observed in our mesocosm study. Although such agregation of bacteria at the phylum level may obscure differences between taxa that exist at finer-scale level (Martiny et al. 2015, Ho et al. 2017), such high level classification shows some ecological coherence and has been key in the developpemnt of general theories in microbial ecology (Philippot et al. 2010). With this consideration in mind, Gram negative bacteria were associated with r strategists bacteria that use labile C to sustain fast growth (De Vries and Shade 2013, Fanin et al. 2018). Such growth strategy can promote microbial biomass and enzymatic processes (Piton et al. 2020b), and eventually the high SMF that we observed. Altogether, a large proportion (25%) of SMF was explained by these four properties (soil moisture, C:N imbalance, bacterial diversity and GP:GN) stressing their coupling in the soil system. However, about the same proportion (24%) was additionally explained only by the microbial properties (bacterial diversity and GP:GN), suggesting that high SMF not only emerged from favorable stoichiometric and soil moisture conditions for microbial growth, but also relied on microbial diversity (Graham et al. 2016).

By examining the combined effects of plant presence, fertilization and drought on soil stoichiometry and microbial diversity, we identified key mechanisms driving SMF in mountain grassland soils. When plants were present, nutrient and water fluxes became tightly coupled with significant consequences for microbial communities (De Vries et al. 2018). Plant growth was associated with an uptake of nutrient and water, which was compensated by water supply and nutrient mineralization in the control treatment. Plants also supplied labile C to the soil through rhizodeposition (Pausch and Kuzyakov 2018), balancing microbial resource stoichiometry, limiting C limitation and promoting microbial biomass and activity, thereby enhancing SMF. However, fertilization induced a massive increase of available N that strengthened C-limitation for microbes but also promoted plant growth, and associated water and nutrient uptake. Without water supply in the drought treatment, the fast growth of plants in the fertilized treatment rapidly decreased soil water content. Below a certain threshold of water availability, plant growth was reduced (Colman and Lazenby 1975, Huang et al. 2020) and root absorption of fertilizer-N stopped. The plant growth under drought were thus not able any more to balance the microbial C limitation through C-rhizodeposition and N-uptake. The combined effects of low moisture and high C limitation induced a decrease of bacteria diversity and selected for gram positive bacteria, as previously reported (Fanin et al. 2018, Zhou et al. 2020), with detrimental consequences for SMF. In unplanted mountain grassland soils, drought induced a weaker effect on soil moisture, soil nutrient stoichiometry and bacteria community, which eventually limited the detrimental drought-related consequences for SMF. Moreover, we observed that microbial biomass in unplanted soils adapted their stoichiometry in response to increased N inputs provided by fertilization, potentially reflecting an accumulation of osmolytes in response to desiccation stress (Schimel et al. 2007). This response of the microbial biomass composition tracked the change of soil resource stoichiometry and limited stoichiometric imbalance and consequences for SMF, a response that we did observed in the planted mountain grassland soil. Altogether, our study showed that combining stoichiometric, life history strategy and biodiversity-function theories represents a powerful approach to understand the mechanisms underlying the complex responses of SMF to global change factors.

## Conclusion

Our study demonstrates that in a mountain grassland system, mineral fertilization with the aim to promote plant productivity could impair the drought resistance of soil multifunctionality. This effect was driven by plant-mediated changes in soil resources availability and microbial diversity which both underlie soil multifunctionality. These findings highlight the importance of integrating plant-soil-microorganisms interactions to fully capture ecosystem responses in global change.

## ACKNOWLEDGEMENTS

We are grateful to Lucas Paysan for providing soil from his grassland and to the Station Alpine Joseph Fourier for their greenhouse. We also thank Cindy Arnoldi, Bastien Audemard, Jonathan Gervaix, Christian Miquel, Viet Tran-Khac for their help in the laboratory work.
Preprint version 2 of this article has been peer-reviewed and recommended by Peer Community In Ecology (https://doi.org/10.24072/pci.ecology.100761; Barot, 2025).

## DATA, SCRIPTS, CODE, AND SUPPLEMENTARY INFORMATION AVAILABILITY

All data and R scripts used in this study are deposited with folowing DOI : 10.5281/zenodo.14962600

## Supplementary materials

### Supplementary figures

**Figure S1.**
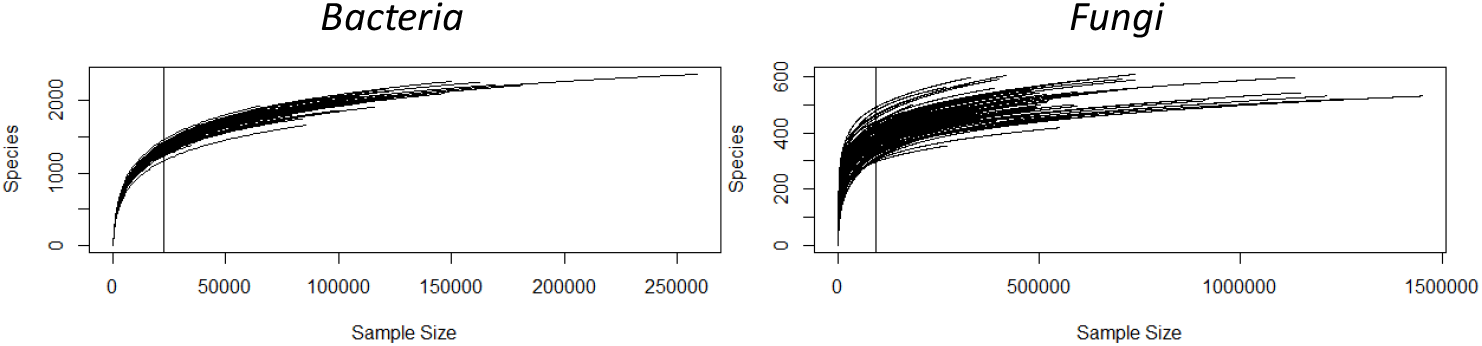
Rarefaction curves for Bacteria and Fungi. Vertical lines represent resampling size used for data rarefaction of each sample to have similar sequencing depth equal to the number of reads of the sample with the lowest reads numbers (Bacteria: 22 571 sequences/sample, Eukaryota : 93 742 sequences/sample).

**Figure S2.**
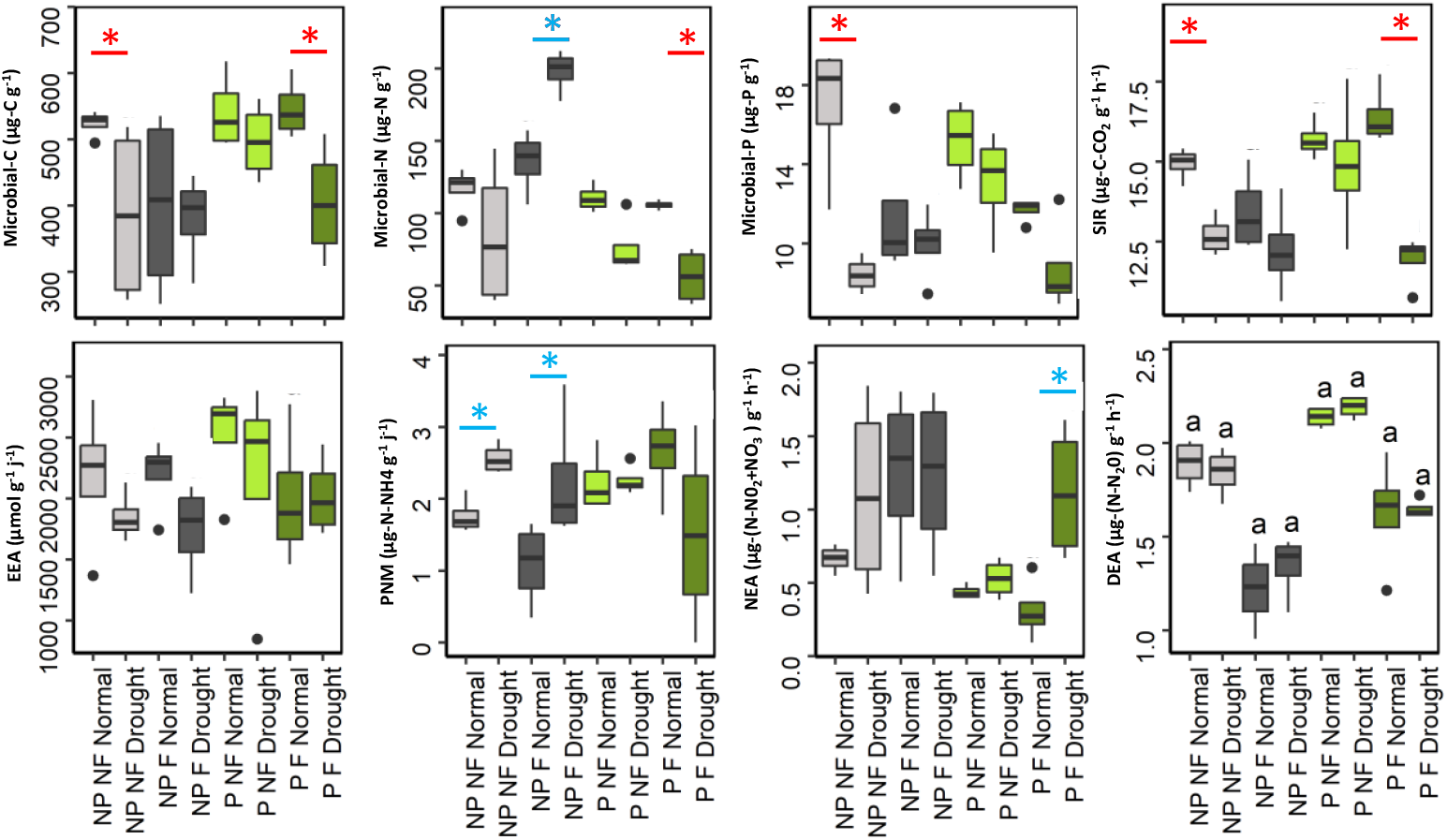
Effect of drought, fertilization and plant presence on soil microbial functions. Star indicate significant difference between drought and normal climate treatment. SIR=Substrate-induced-respiration. PNM=Potential N mineralization. NEA=Potential nitrification activity. DEA=Potential denitrification activity. NP=No plant, P=Plant presence, F=Fertilized, NF=No fertilization

**Figure S3.**
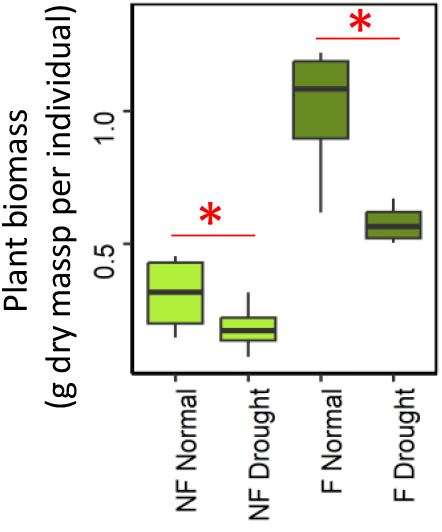
Plant biomass response to drought with or without fertilisation. Star indicate significant difference between drought and normal climate treatment.

**Figure S4.**
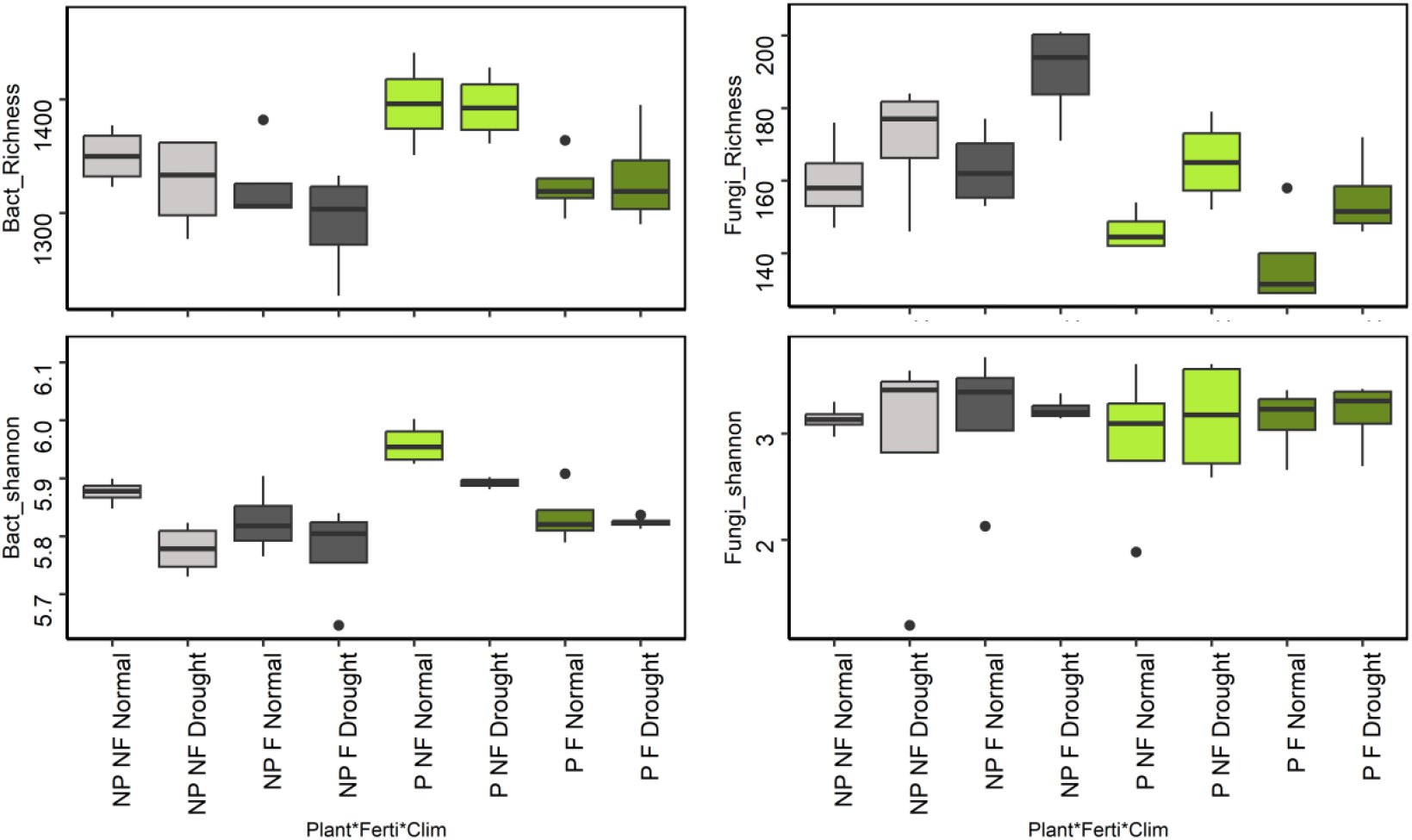
Soil microbial diversity response to drought, fertilization and plant presence. Model statistics are provided in Table S2. NP=No plant, P=Plant presence, F=Fertilized, NF=No fertilization

**Figure S5.**
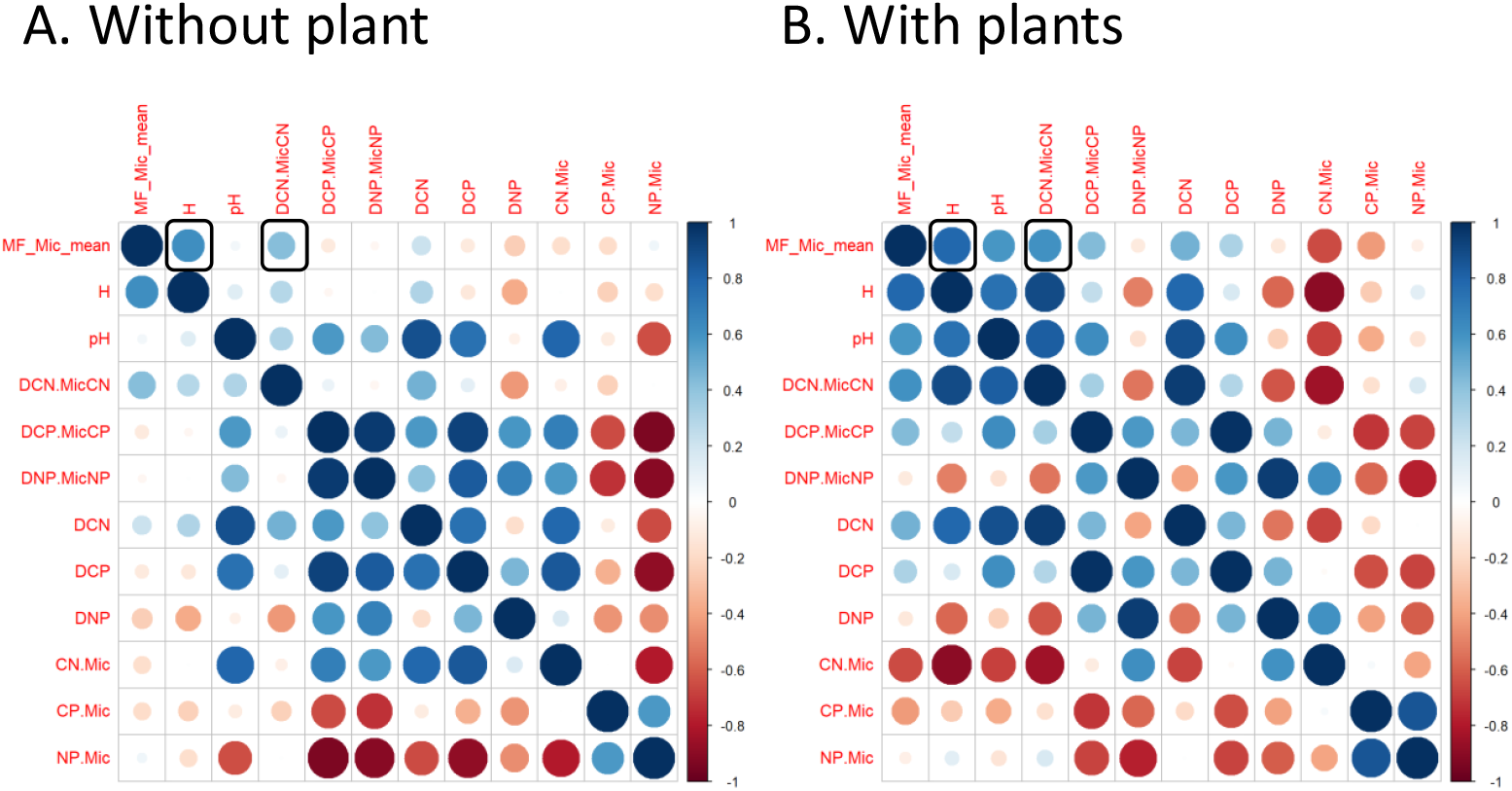
Spearman correlation matrix between soil multifunctionality (MF_Mic_mean), soil moisture, pH and stoichiometry (DCN=DOC:TDN, DCP=DOC:PO4, DNP=TDN:PO4) in soil without plants (A) or with plants (B). Circle color represent correlation direction and circle size and color intensity represent correlation coefficient.

**Figure S6.**
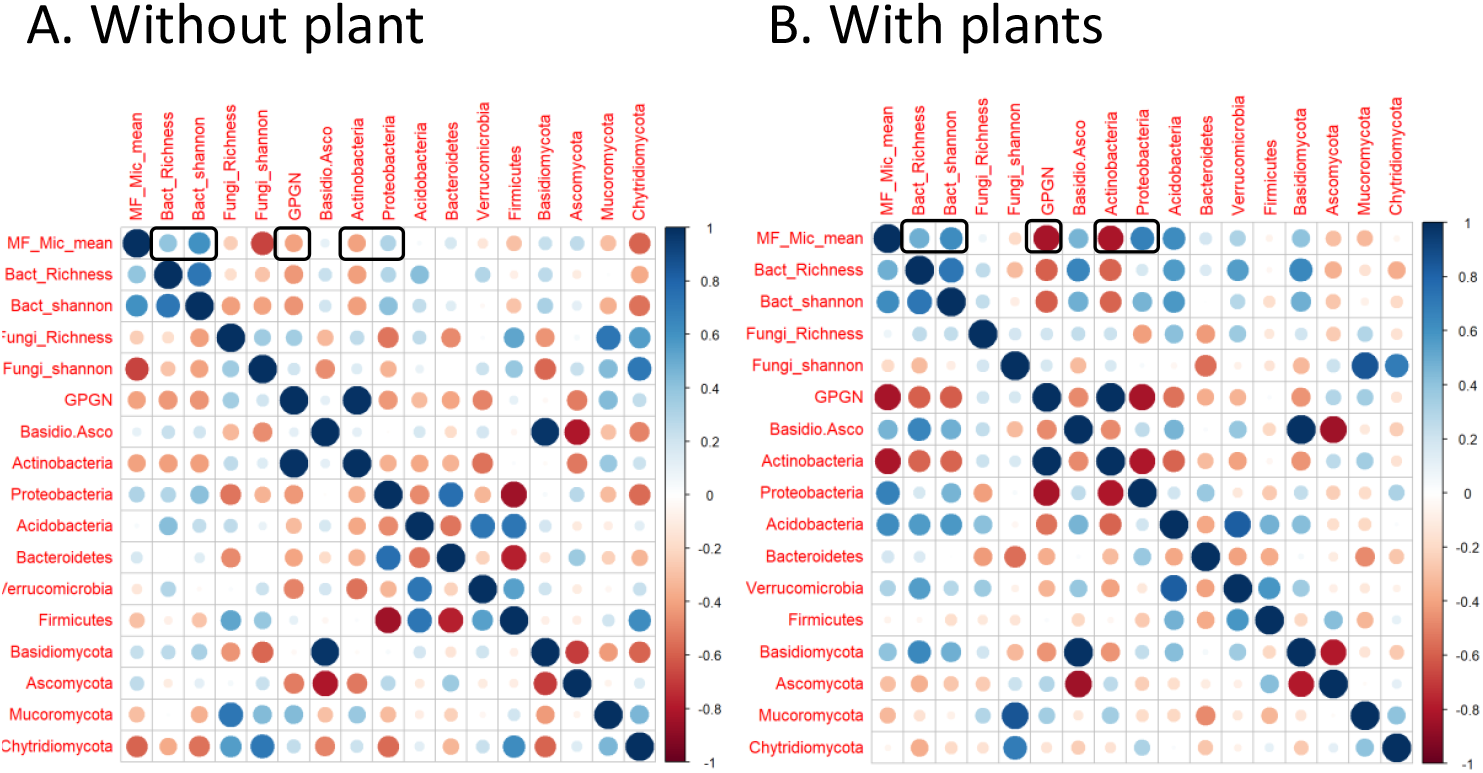
Spearman correlation matrix between soil multifunctionality (MF_Mic_mean), soil microbial community diversity, phylum relative abundances and gram positive : gram negative ratio (GPGN) in soil without plant (A) or with plants (B). Circle color represent correlation direction and circle size and color intensity represent correlation coefficient. Cells in black frames are consistent relationship across soil with and without plants.

**Figure S7.**
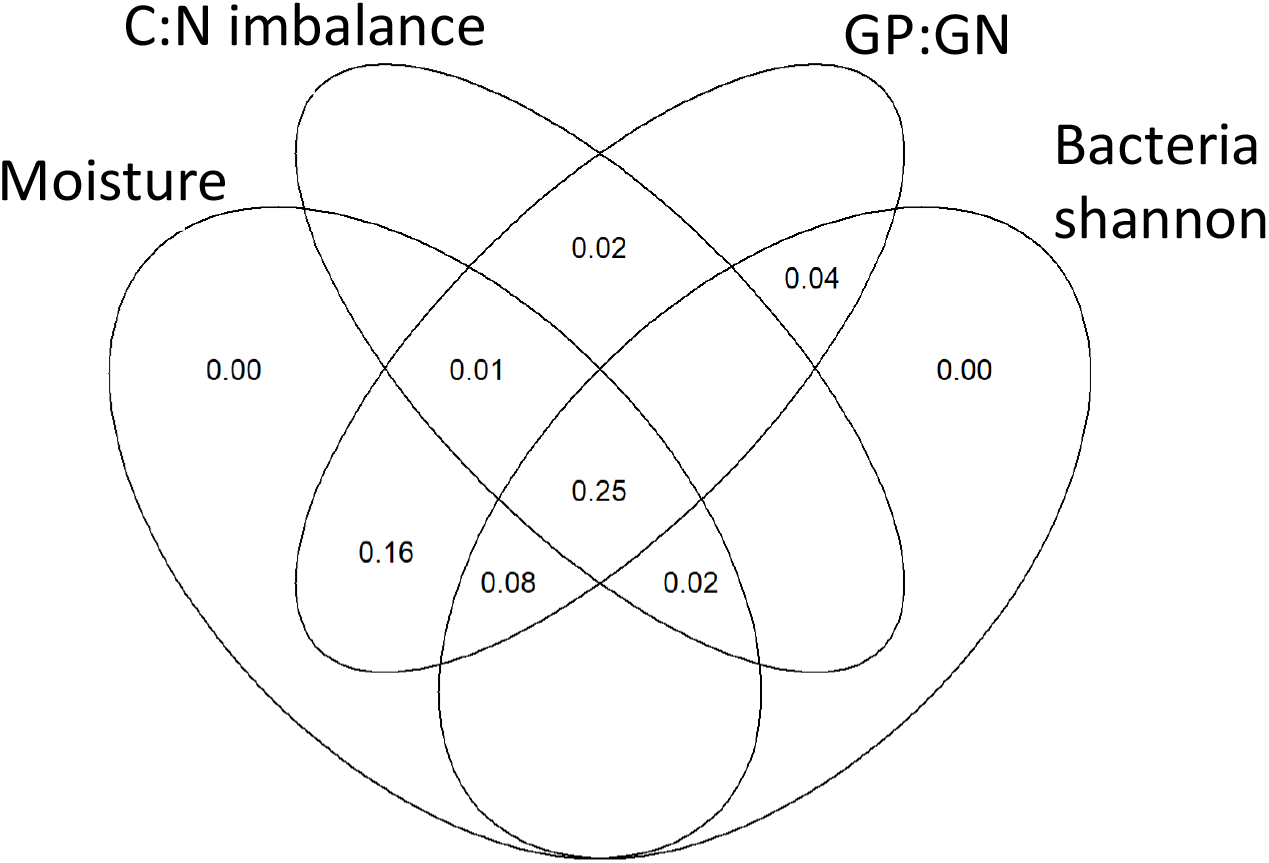
Venn diagram of variance partitioning analysis illustrating the effect of moisture, C:N imbalance, GP:GN ratio, and Bacterial Shannon index on SMF. Values show the percentage of explained variance by each variable, and the percentage commonly explained in the intersections.

#### Supplementary Tables

**Table S1.**
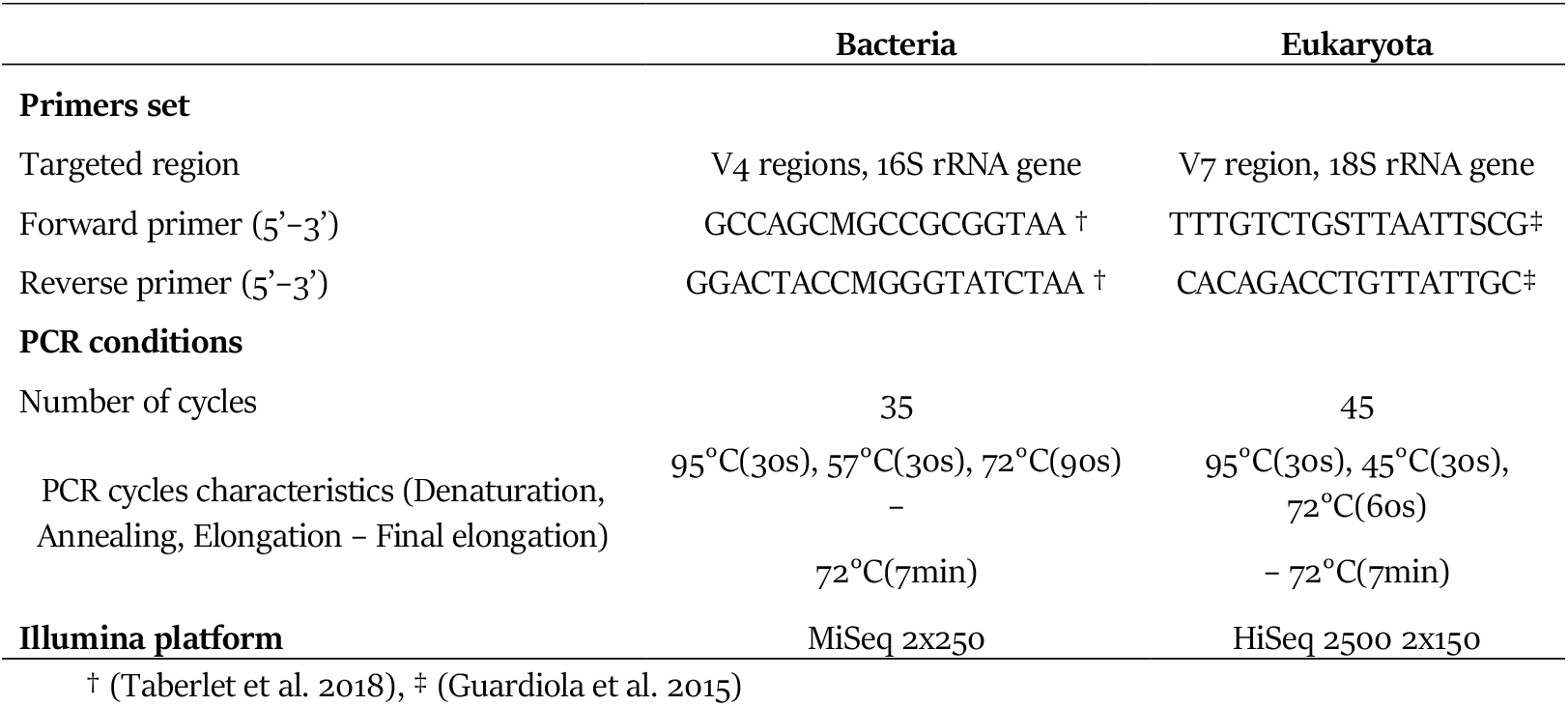
Overview of primers characteristics and cycling conditions used for polymerase chain reaction (PCR) and amplicon sequencing.

**Table S2.**
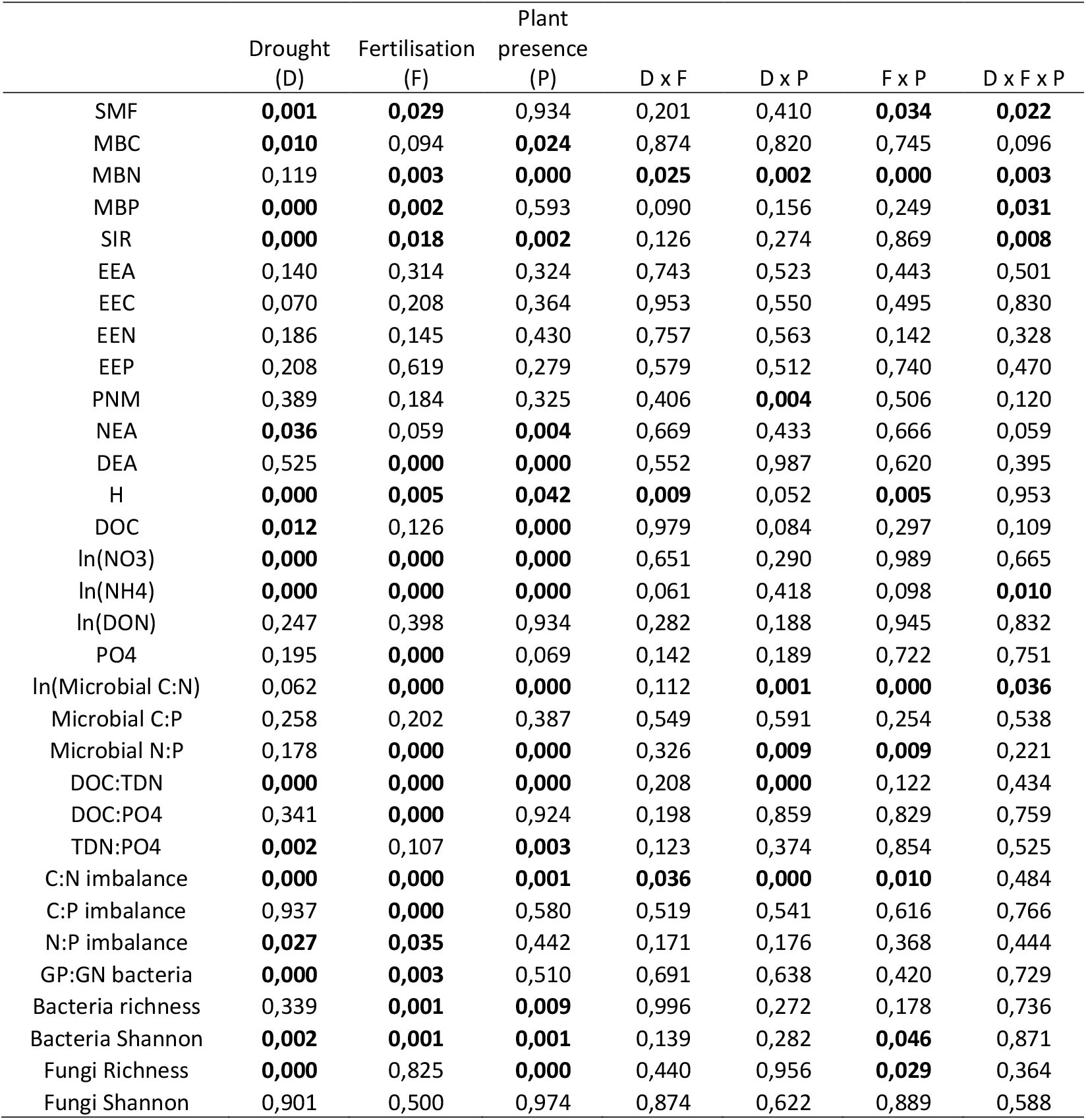
P-values of ANOVA 3 testing the effect of Drought, Fertilization, Plant presence and their interactions in SMF, microbial functions, soil chemistry, stoechiometry and microbial diversity. SMF=Soil multifunctionality, MBC=Microbial biomass-C, MBN=Microbial biomass-N, MBP=Microbial biomass-P, SIR=Substrate induced respiration, EEA= cumulative enzyme activity, EEC=C-acquisition enzyme activity, EEN= N-acquisition enzyme activity, EEP=P-acquisition enzyme activity, PNM=Potential Nitrogen mineralization, NEA=Potential nitrification enzyme activity, DEA= Potential denitrification activity, H=soil humidity, DOC=Dissolved organic C, NO3=Nitraten NH4=Amonium, DON=Dissolved organic N, PO4=Phosphate, GP:GN=ratio of Gram positive : Gram negative bacteria, Shannon= Shannon diversity index.

## References

Allison, S. D., and J. B. Martiny. 2008. Resistance, resilience, and redundancy in microbial communities. Proceedings of the National Academy of Sciences 105:11512–11519.

Anderson, J., and K. Domsch. 1978. A physiological method for the quantitative measurement of microbial biomass in soils. Soil biology and biochemistry 10:215–221.

Attard, E., S. Recous, A. Chabbi, C. De Berranger, N. Guillaumaud, J. Labreuche, L. Philippot, B. Schmid, and X. Le Roux. 2011. Soil environmental conditions rather than denitrifier abundance and diversity drive potential denitrification after changes in land uses. Global Change Biology 17:1975–1989.

Bardgett, R. D., J. M. Bullock, S. Lavorel, P. Manning, U. Schaffner, N. Ostle, M. Chomel, G. Durigan, E. L Fry, D. Johnson, and others. 2021. Combatting global grassland degradation. Nature Reviews Earth & Environment 2:720–735.

Bardgett, R. D., C. Freeman, and N. J. Ostle. 2008. Microbial contributions to climate change through carbon cycle feedbacks. The ISME journal 2:805–814.

Bardgett, R. D., P. J. Hobbs, and Å. Frostegård. 1996. Changes in soil fungal: bacterial biomass ratios following reductions in the intensity of management of an upland grassland. Biology and Fertility of Soils 22:261– 264.

Bell, C. W., B. E. Fricks, J. D. Rocca, J. M. Steinweg, S. K. McMahon, and M. D. Wallenstein. 2013. High-throughput fluorometric measurement of potential soil extracellular enzyme activities. Journal of visualized experiments: JoVE.

Bloor, J. M., M. Zwicke, and C. Picon-Cochard. 2018. Drought responses of root biomass provide an indicator of soil microbial drought resistance in grass monocultures. Applied Soil Ecology 126:160–164.

Borken, W., and E. Matzner. 2009. Reappraisal of drying and wetting effects on C and N mineralization and fluxes in soils. Global Change Biology 15:808–824.

Boyer, F., C. Mercier, A. Bonin, Y. Le Bras, P. Taberlet, and E. Coissac. 2016. obitools: A unix-inspired software package for DNA metabarcoding. Molecular ecology resources 16:176–182.

Brookes, P., D. Powlson, and D. Jenkinson. 1982. Measurement of microbial biomass phosphorus in soil. Soil biology and biochemistry 14:319–329.

Brunner, M. I., A. F. Van Loon, and K. Stahl. 2022. Moderate and severe hydrological droughts in Europe differ in their hydrometeorological drivers. Water Resources Research 58:e2022WR032871.

Camenzind, T., C. A. Aguilar-Trigueros, S. Hempel, A. Lehmann, M. Bielcik, D. R. Andrade-Linares, J. Bergmann, J. dela Cruz, J. Gawronski, P. Golubeva, and others. 2024. Towards establishing a fungal economics spectrum in soil saprobic fungi. Nature Communications 15:3321.

Canarini, A., L. P. Kiær, and F. A. Dijkstra. 2017. Soil carbon loss regulated by drought intensity and available substrate: A meta-analysis. Soil Biology and Biochemistry 112:90–99.

Colman, R., and A. Lazenby. 1975. Effect of moisture on growth and nitrogen response by Lolium perenne. Plant and Soil 42:1–13.

Crowther, T. W., J. Van den Hoogen, J. Wan, M. A. Mayes, A. Keiser, L. Mo, C. Averill, and D. S. Maynard. 2019. The global soil community and its influence on biogeochemistry. Science 365:eaav0550.

Dassonville, N., N. Guillaumaud, F. Piola, P. Meerts, and F. Poly. 2011. Niche construction by the invasive Asian knotweeds (species complex Fallopia): impact on activity, abundance and community structure of denitrifiers and nitrifiers. Biological invasions 13:1115–1133.

Delgado-Baquerizo, M., D. J. Eldridge, V. Ochoa, B. Gozalo, B. K. Singh, and F. T. Maestre. 2017. Soil microbial communities drive the resistance of ecosystem multifunctionality to global change in drylands across the globe. Ecology letters 20:1295–1305.

Delgado-Baquerizo, M., L. Giaramida, P. B. Reich, A. N. Khachane, K. Hamonts, C. Edwards, L. A. Lawton, and B. K. Singh. 2016a. Lack of functional redundancy in the relationship between microbial diversity and ecosystem functioning. Journal of ecology 104:936–946.

Delgado-Baquerizo, M., F. T. Maestre, P. B. Reich, T. C. Jeffries, J. J. Gaitan, D. Encinar, M. Berdugo, C. D. Campbell, and B. K. Singh. 2016b. Microbial diversity drives multifunctionality in terrestrial ecosystems. Nature communications 7:10541.

Dong, L., X. Yao, Y. Deng, H. Zhang, W. Zeng, X. Li, J. Tang, and W. Wang. 2022. Nitrogen deficiency in soil mediates multifunctionality responses to global climatic drivers. Science of the Total Environment 838:156533.

Eskelinen, A., K. Gravuer, W. S. Harpole, S. Harrison, R. Virtanen, and Y. Hautier. 2020. Resource-enhancing global changes drive a whole-ecosystem shift to faster cycling but decrease diversity. Ecology 101:e03178.

Evans, S. E., and M. D. Wallenstein. 2014. Climate change alters ecological strategies of soil bacteria. Ecology letters 17:155–164.

Fanin, N., P. Kardol, M. Farrell, M.-C. Nilsson, M. J. Gundale, and D. A. Wardle. 2018. The ratio of Gram-positive to Gram-negative bacterial PLFA markers as an indicator of carbon availability in organic soils. Soil Biology and Biochemistry.

Fierer, N. 2017. Embracing the unknown: disentangling the complexities of the soil microbiome. Nature Reviews Microbiology 15:579–590.

Fierer, N., M. A. Bradford, and R. B. Jackson. 2007. Toward an ecological classification of soil bacteria. Ecology 88:1354–1364.

Garland, G., S. Banerjee, A. Edlinger, E. Miranda Oliveira, C. Herzog, R. Wittwer, L. Philippot, F. T. Maestre, and M. G. van Der Heijden. 2021. A closer look at the functions behind ecosystem multifunctionality: A review. Journal of Ecology 109:600–613.

Graham, E. B., J. E. Knelman, A. Schindlbacher, S. Siciliano, M. Breulmann, A. Yannarell, J. Beman, G. Abell, L. Philippot, J. Prosser, and others. 2016. Microbes as engines of ecosystem function: when does community structure enhance predictions of ecosystem processes? Frontiers in microbiology 7:214.

Grigulis, K., S. Lavorel, U. Krainer, N. Legay, C. Baxendale, M. Dumont, E. Kastl, C. Arnoldi, R. D. Bardgett, F. Poly, and others. 2013. Relative contributions of plant traits and soil microbial properties to mountain grassland ecosystem services. Journal of Ecology 101:47–57.

Guardiola, M., M. J. Uriz, P. Taberlet, E. Coissac, O. S. Wangensteen, and X. Turon. 2015. Deep-sea, deep-sequencing: metabarcoding extracellular DNA from sediments of marine canyons. PLoS One 10:e0139633.

Gupta, R. S. 2011. Origin of diderm (Gram-negative) bacteria: antibiotic selection pressure rather than endosymbiosis likely led to the evolution of bacterial cells with two membranes. Antonie Van Leeuwenhoek 100:171–182.

Ho, A., D. P. Di Lonardo, and P. L. Bodelier. 2017. Revisiting life strategy concepts in environmental microbial ecology. FEMS microbiology ecology 93:fix006.

Huang, Z., Y. Liu, F.-P. Tian, and G.-L. Wu. 2020. Soil water availability threshold indicator was determined by using plant physiological responses under drought conditions. Ecological Indicators 118:106740.

Jenkinson, D. S., P. C. Brookes, and D. S. Powlson. 2004. Measuring soil microbial biomass. Soil biology and biochemistry 36:5–7.

Jia, J., G. Hu, G. Ni, M. Xie, R. Li, G. Wang, and J. Zhang. 2024. Bacteria drive soil multifunctionality while fungi are effective only at low pathogen abundance. Science of the Total Environment 906:167596.

Jia, J., J. Zhang, Y. Li, M. Xie, G. Wang, and J. Zhang. 2022. Land use intensity constrains the positive relationship between soil microbial diversity and multifunctionality. Plant and Soil:1–14.

Karlowsky, S., A. Augusti, J. Ingrisch, R. Hasibeder, M. Lange, S. Lavorel, M. Bahn, and G. Gleixner. 2018. Land use in mountain grasslands alters drought response and recovery of carbon allocation and plant-microbial interactions. Journal of Ecology 106:1230–1243.

Koper, T. E., J. M. Stark, M. Y. Habteselassie, and J. M. Norton. 2010. Nitrification exhibits Haldane kinetics in an agricultural soil treated with ammonium sulfate or dairy-waste compost. FEMS microbiology ecology 74:316–322.

Koyama, A., J. M. Steinweg, M. L. Haddix, J. S. Dukes, and M. D. Wallenstein. 2017. Soil bacterial community responses to altered precipitation and temperature regimes in an old field grassland are mediated by plants. FEMS microbiology ecology 94:fix156.

Legay, N., G. Piton, C. Arnoldi, L. Bernard, M.-N. Binet, B. Mouhamadou, T. Pommier, S. Lavorel, A. Foulquier, and J.-C. Clément. 2017. Soil legacy effects of climatic stress, management and plant functional composition on microbial communities influence the response of Lolium perenne to a new drought event. Plant and Soil:1–22.

Lineweaver, H., and D. Burk. 1934. The determination of enzyme dissociation constants. Journal of the American chemical society 56:658–666.

Loucougaray, G., L. Dobremez, P. Gos, Y. Pauthenet, B. Nettier, and S. Lavorel. 2015. Assessing the effects of grassland management on forage production and environmental quality to identify paths to ecological intensification in mountain grasslands. Environmental management 56:1039–1052.

Luo, G., T. Wang, K. Li, L. Li, J. Zhang, S. Guo, N. Ling, and Q. Shen. 2019. Historical nitrogen deposition and straw addition facilitate the resistance of soil multifunctionality to drying-wetting cycles. Applied and Environmental Microbiology 85:2251–18.

Maestre, F. T., J. L. Quero, N. J. Gotelli, A. Escudero, V. Ochoa, M. Delgado-Baquerizo, M. Garcia-Gómez, M. A. Bowker, S. Soliveres, C. Escolar, and others. 2012. Plant species richness and ecosystem multifunctionality in global drylands. Science 335:214–218.

Manzoni, S., M. H. Ahmed, and A. Porporato. 2019. Ecohydrological and stoichiometric controls on soil carbon and nitrogen dynamics in drylands. Dryland Ecohydrology:279–307.

Maron, P.-A., A. Sarr, A. Kaisermann, J. Lévêque, O. Mathieu, J. Guigue, B. Karimi, L. Bernard, S. Dequiedt, S. Terrat, and others. 2018. High microbial diversity promotes soil ecosystem functioning. Appl. Environ. Microbiol. 84:2738–17.

Martiny, J. B., S. E. Jones, J. T. Lennon, and A. C. Martiny. 2015. Microbiomes in light of traits: a phylogenetic perspective. Science 350:aac9323.

Mercier, C., F. Boyer, A. Bonin, and E. Coissac. 2013. SUMATRA and SUMACLUST: fast and exact comparison and clustering of sequences. Pages 27–29 Programs and Abstracts of the SeqBio 2013 workshop. Abstract. . Citeseer.

Möhl, P., M. Vorkauf, A. Kahmen, and E. Hiltbrunner. 2023. Recurrent summer drought affects biomass production and community composition independently of snowmelt manipulation in alpine grassland. Journal of Ecology 111:2357–2375.

Mooshammer, M., W. Wanek, S. Zechmeister-Boltenstern, and A. A. Richter. 2014. Stoichiometric imbalances between terrestrial decomposer communities and their resources: mechanisms and implications of microbial adaptations to their resources. Frontiers in microbiology 5:22.

Murphy, J., and J. P. Riley. 1962. A modified single solution method for the determination of phosphate in natural waters. Analytica chimica acta 27:31–36.

Naylor, D., and D. Coleman-Derr. 2018. Drought stress and root-associated bacterial communities. Frontiers in plant science 8:2223.

Oksanen, J., F. G. Blanchet, R. Kindt, P. Legendre, P. R. Minchin, R. O’hara, G. L. Simpson, P. Solymos, M. H. H. Stevens, and H. Wagner. 2011. vegan: Community ecology package. R package version:117– 118.

Orwin, K., I. Dickie, R. Holdaway, and J. Wood. 2018. A comparison of the ability of PLFA and 16S rRNA gene metabarcoding to resolve soil community change and predict ecosystem functions. Soil Biology and Biochemistry 117:27–35.

Orwin, K. H., and D. A. Wardle. 2005. Plant species composition effects on belowground properties and the resistance and resilience of the soil microflora to a drying disturbance. Plant and Soil 278:205–221.

Osburn, E. D., G. Yang, M. C. Rillig, and M. S. Strickland. 2023. Evaluating the role of bacterial diversity in supporting soil ecosystem functions under anthropogenic stress. ISME communications 3:66.

Pausch, J., and Y. Kuzyakov. 2018. Carbon input by roots into the soil: quantification of rhizodeposition from root to ecosystem scale. Global change biology 24:1–12.

Philippot, L., S. G. Andersson, T. J. Battin, J. I. Prosser, J. P. Schimel, W. B. Whitman, and S. Hallin. 2010. The ecological coherence of high bacterial taxonomic ranks. Nature Reviews Microbiology 8:523–529.

Piton, G., S. D. Allison, M. Bahram, F. Hildebrand, J. B. Martiny, K. K. Treseder, and A. C. Martiny. 2023. Life history strategies of soil bacterial communities across global terrestrial biomes. Nature microbiology:1–10.

Piton, G., A. Foulquier, L. B. Martinez-Garcia, N. Legay, C. Arnoldi, L. Brussaard, K. Hedlund, P. Martins da Silva, E. Nascimento, F. Reis, and others. 2020a. Resistance–recovery trade-off of soil microbial communities under altered rain regimes: An experimental test across European agroecosystems. Journal of Applied Ecology.

Piton, G., A. Foulquier, L. B. Martinez-Garcia, N. Legay, K. Hedlund, P. M. da Silva, E. Nascimento, F. Reis, P. Sousa, G. B. De Deyn, and others. 2020b. Disentangling drivers of soil microbial potential enzyme activity across rain regimes: An approach based on the functional trait framework. Soil Biology and Biochemistry:107881.

Piton, G., N. Legay, C. Arnoldi, S. Lavorel, J. C. Clément, and A. Foulquier. 2020c. Using proxies of microbial community-weighted means traits to explain the cascading effect of management intensity, soil and plant traits on ecosystem resilience in mountain grasslands. Journal of Ecology 108:876–893.

Preece, C., G. Farré-Armengol, and J. Peñuelas. 2020. Drought is a stronger driver of soil respiration and microbial communities than nitrogen or phosphorus addition in two Mediterranean tree species. Science of The Total Environment 735:139554.

Robertson, G. P., D. C. Coleman, P. Sollins, C. S. Bledsoe, and others. 1999. Standard soil methods for long-term ecological research. Oxford University Press on Demand.

Schimel, J., T. C. Balser, and M. Wallenstein. 2007. Microbial stress-response physiology and its implications for ecosystem function. Ecology 88:1386–1394.

Schnell, I. B., K. Bohmann, and M. T. P. Gilbert. 2015. Tag jumps illuminated–reducing sequence-to-sample misidentifications in metabarcoding studies. Molecular ecology resources 15:1289–1303.

Shukla, P., J. Skea, E. Calvo Buendia, V. Masson-Delmotte, H. Pörtner, D. Roberts, P. Zhai, R. Slade, S. Connors, R. Van Diemen, and others. 2019. IPCC, 2019: Climate Change and Land: an IPCC special report on climate change, desertification, land degradation, sustainable land management, food security, and greenhouse gas fluxes in terrestrial ecosystems.

Taberlet, P., A. Bonin, L. Zinger, and E. Coissac. 2018. Environmental DNA: For biodiversity research and monitoring. Oxford University Press.

Taberlet, P., E. Coissac, F. Pompanon, C. Brochmann, and E. Willerslev. 2012. Towards next-generation biodiversity assessment using DNA metabarcoding. Molecular ecology 21:2045–2050.

Tardy, V., O. Mathieu, J. Lévêque, S. Terrat, A. Chabbi, P. Lemanceau, L. Ranjard, and P.-A. Maron. 2014. Stability of soil microbial structure and activity depends on microbial diversity. Environmental microbiology reports 6:173–183.

Trivedi, C., M. Delgado-Baquerizo, K. Hamonts, K. Lai, P. B. Reich, and B. K. Singh. 2019. Losses in microbial functional diversity reduce the rate of key soil processes. Soil Biology and Biochemistry 135:267–274.

Vance, E. D., P. C. Brookes, and D. S. Jenkinson. 1987. An extraction method for measuring soil microbial biomass C. Soil biology and Biochemistry 19:703–707.

De Vries, F. T., and R. D. Bardgett. 2012. Plant-microbial linkages and ecosystem nitrogen retention: lessons for sustainable agriculture. Frontiers in Ecology and the Environment 10:425–432.

De Vries, F. T., R. I. Griffiths, M. Bailey, H. Craig, M. Girlanda, H. S. Gweon, S. Hallin, A. Kaisermann, A. M. Keith, M. Kretzschmar, and others. 2018. Soil bacterial networks are less stable under drought than fungal networks. Nature communications 9:3033.

De Vries, F. T., M. E. Liiri, L. Bjørnlund, M. A. Bowker, S. Christensen, H.M. Setälä, and R. D. Bardgett. 2012. Land use alters the resistance and resilience of soil food webs to drought. Nature Climate Change 2:276–280.

De Vries, F. T., and A. Shade. 2013. Controls on soil microbial community stability under climate change. Frontiers in microbiology 4:265.

Wardle, D. A., and M. Jonsson. 2014. Long-term resilience of above-and belowground ecosystem components among contrasting ecosystems. Ecology 95:1836–1849.

Wienhold, B. J. 2007. Comparison of laboratory methods and an in situ method for estimating nitrogen mineralization in an irrigated silt-loam soil. Communications in soil science and plant analysis 38:1721– 1732.

Yachi, S., and M. Loreau. 1999. Biodiversity and ecosystem productivity in a fluctuating environment: the insurance hypothesis. Proceedings of the National Academy of Sciences 96:1463–1468.

Yang, G., M. Ryo, J. Roy, D. R. Lammel, M.-B. Ballhausen, X. Jing, X. Zhu, and M. C. Rillig. 2022. Multiple anthropogenic pressures eliminate the effects of soil microbial diversity on ecosystem functions in experimental microcosms. Nature communications 13:4260.

Yang, X., X. Huang, Z. Cheng, S. Li, H. Mahjoob, J. Xu, and Y. He. 2024. Response of soil general and specific functions following loss of microbial diversity: A review. Soil Security:100151.

Zavaleta, E. S., J. R. Pasari, K. B. Hulvey, and G. D. Tilman. 2010. Sustaining multiple ecosystem functions in grassland communities requires higher biodiversity. Proceedings of the National Academy of Sciences 107:1443–1446.

Zechmeister-Boltenstern, S., K. M. Keiblinger, M. Mooshammer, J. Peñuelas, A. Richter, J. Sardans, and W. Wanek. 2015. The application of ecological stoichiometry to plant–microbial–soil organic matter transformations. Ecological Monographs 85:133–155.

Zhou, Z., C. Wang, and Y. Luo. 2020. Meta-analysis of the impacts of global change factors on soil microbial diversity and functionality. Nature communications 11:3072.

Zinger, L., P. Taberlet, H. Schimann, A. Bonin, F. Boyer, M. De Barba, P. Gaucher, L. Gielly, C. Giguet-Covex, A. Iribar, and others. 2019. Body size determines soil community assembly in a tropical forest. Molecular ecology 28:528–543.

